# [ICoN: Integration using Co-attention across Biological Networks

**DOI:** 10.1101/2024.02.05.577786

**Authors:** Nure Tasnina, T. M. Murali

**Affiliations:** Dept. of Computer Science, Virginia Tech, Blacksburg, VA 24061, USA

**Keywords:** network integration, graph neural networks, co-attention

## Abstract

**Motivation:** Molecular interaction networks are powerful tools for studying cellular functions. Integrating diverse types of networks enhances performance in downstream tasks such as gene module detection and protein function prediction. The challenge lies in extracting meaningful protein feature representations due to varying levels of sparsity and noise across these heterogeneous networks.

**Results:** We propose ICoN, a novel ‘co-attention’-based, denoising, unsupervised graph neural network model that takes multiple protein-protein association networks as inputs and generates an integrated single network by computing a unified feature representation for each protein. A key contribution of ICoN is a novel approach that enables cross-network communication through co-attention during training. The model also incorporates a denoising training technique, introducing perturbations to each input network and training the model to reconstruct the original network from its corrupted version, a method previously unexplored in network integration.

Our experimental results demonstrate that ICoN surpasses individual networks across three downstream tasks: gene module detection, gene coannotation prediction, and protein function prediction. Compared to existing unsupervised network integration models, ICoN exhibits superior performance across the majority of downstream tasks and exhibits enhanced robustness against noise. This work introduces a promising approach for effectively integrating diverse protein-protein association networks, aiming to achieve a biologically meaningful unified representation of proteins.

**Availability:** The ICoN software is available under the GNU Public License v3 at https://github.com/Murali-group/ICoN.

## 1 Introduction

The emergence of high-throughput experimental techniques has led to the development of extensive protein-protein association networks. In such a network, each node is a protein and an edge between two nodes represents some type of association, e.g., physical binding, shared cellular function, or correlated expression. In principle, each type of interaction yields a separate network [1, 2]. Since different types of networks provide heterogeneous and complementary biological information, integrating them into a common representation allows for improved performance over using a single data source in tackling questions such as protein function prediction and gene module detection [3, 4]. However, integrating these networks, i.e., combining all the networks into one single network, is not trivial. The integration process poses a significant challenge due to the presence of false positive and false negative edges resulting from experiments with noise and limited resources, respectively. Early techniques for network integration relied on supervision using protein function annotations as labels [5, 6]. However, the performance of these models is contingent on the availability and quality of the provided labels. Moreover, these models may not generalize well to other downstream tasks to which the integrated networks can be applied.

An important aspect of network integration is acquiring a unified feature representation (embedding) for every protein that encompasses topological information from all input networks. Four recent methods, Mashup [7], deepNF [8], BIONIC [4], and BERTWalk [9] have presented unsupervised approaches to generate unified protein embeddings and showed promising performance in terms of downstream analysis, e.g., protein function prediction and module detection. Each of these methods computes initial embeddings independently for each network. These embeddings encode all the topological information for each network. They do not allow this information to be shared across the input networks during training.

We hypothesized that a model that facilitates continuous inter-network communication during training may learn a more unified representation of proteins which in turn would yield superior performance in downstream tasks. In this context, we embraced the notion of multi-modal alignment, as commonly utilized in vision-language models [10, 11]. This approach facilitates integration across diverse data types such as image, text, and video by leveraging shared attention mechanisms across modality-specific transformer encoders [10]. We adapt this idea to the context of graphs by introducing a co-attention mechanism that aligns multiple topological contexts of a single protein originating from heterogeneous networks to obtain a unified embedding for each protein.

In an attempt to make the integration model robust to noise, we adopt a training technique inspired by denoising autoencoders (DA) [12, 13, 14]. In principle, a DA trains a model to learn the original inputs from corrupted or noisy data by deliberately introducing noise into the input during the training process.

### Our contributions

We propose ICoN (“**I**ntegration using **Co**-attention across Biological **N**etworks”), a novel co-attention-based, denoising, unsupervised graph neural network model that takes multiple protein association networks as inputs and generates an integrated single network by computing a unified feature representation for each protein. A traditional GAT model generates a node’s embeddings by aggregating features from its neighbors in only one network, learning varying degrees of priority or attention for each neighbor. In contrast, ICoN learns the attention given to other nodes by considering the topological context of that node across *all* input networks. This concept of co-attention permits cross-network communication during training. Furthermore, we adopt a denoising training technique where we introduce perturbation to each input network and train the model to reconstruct the original one from the corrupted version. To the best of our knowledge, this method has not been previously employed in the context of network integration.

### Our results

We evaluate ICoN using the evaluation framework developed by BIONIC that includes three downstream tasks (gene coannotation prediction, gene module detection, and gene function prediction) and three benchmark datasets. We demonstrate that the embeddings produced by ICoN outperform those of individual networks across all downstream tasks. In the majority of downstream tasks, ICoN outranked existing unsupervised network integration models. We also demonstrate that the co-attention coefficients learned by ICoN can be interpreted meaningfully in the context of the noise we artificially add during training. Furthermore, we illustrate ICoN’s superior robustness towards noise compared to BIONIC.

## 2 Algorithms

For each network, we consider the union of the nodes across all the input networks, i.e., a node not originally in a network will now appear as disconnected in this network. The starting feature for every node is a onehot encoding, i.e., a vector whose length is equal to the total number of nodes and the index denoting the corresponding node contains 1 while the rest of the indices have the value 0.

### 2.1 Model architecture

There are three principal blocks in ICoN (Figure 1): (i) the **noise induction module** that introduces a specific amount of noise to each of the input networks before inputting them to the subsequent encoder module. (ii) the **Encoder module** computes the embedding of each protein allowing cross-network communication among input networks. (iii) the **network reconstruction module** constructs an integrated network after learning a unified embedding for each protein.

**Figure 1:**
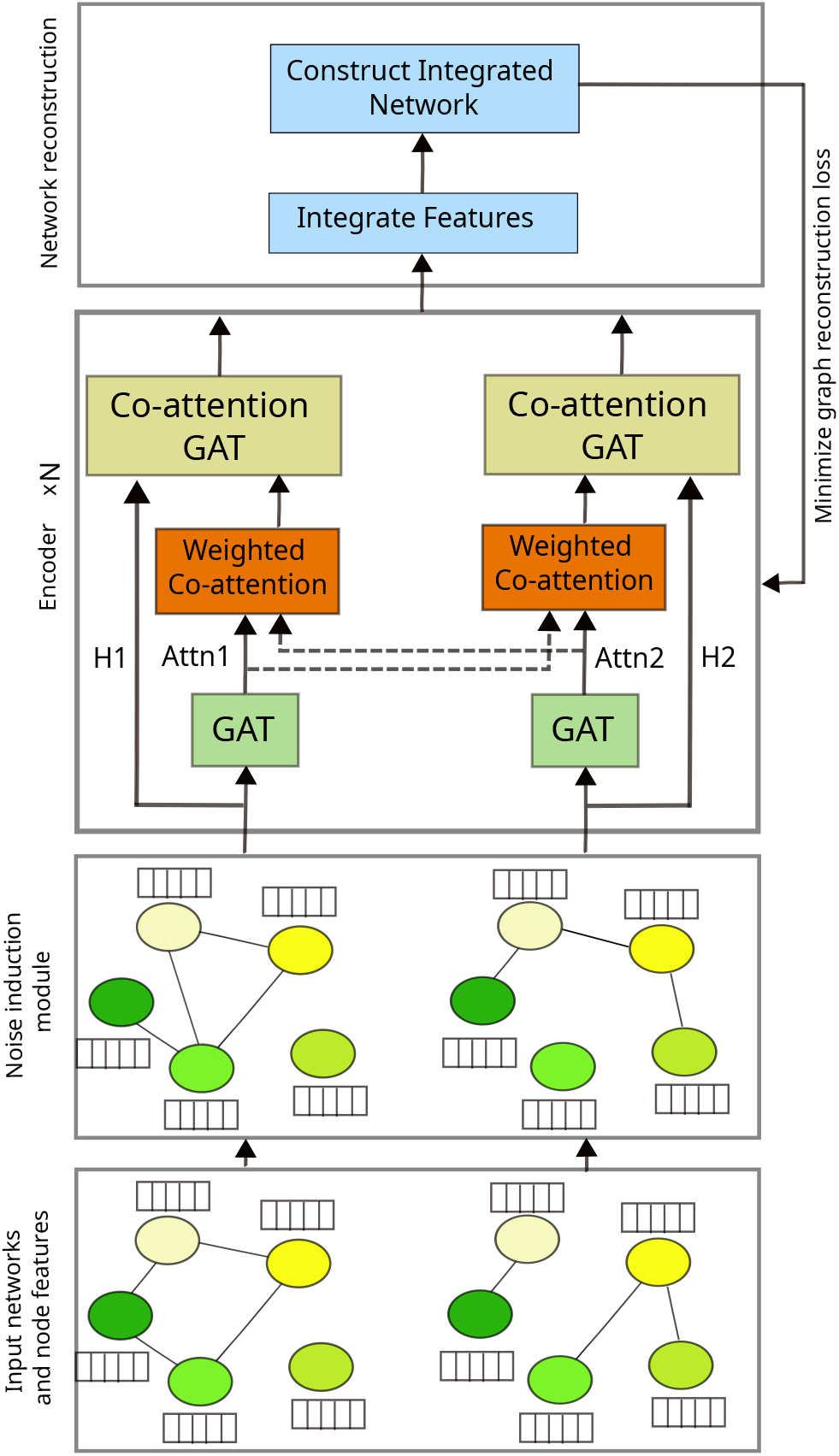
Architecture of ICoN. Attn1 and Attn2 are the learned attention matrices from the corresponding GAT layers.

#### Noise induction module

We introduce noise to each of the input networks by randomly removing a fraction (dictated by hyperparameter *p*) of existing edges and incorporating the same number of new edges, selected uniformly at random from the set of all node pairs not connected by an edge in the network.

#### Encoder module

This module consists of multiple (*N*) stacked blocks of the same architecture. Each block has two main components: (i) a traditional GAT layer that computes the attention of each node towards its neighboring nodes and (ii) a novel co-attention-based GAT that incorporates the attention computed by the first GAT layer for individual input networks and utilizes that to learn a cross-network aware representation of each protein.

#### GAT

Given the adjacency matrix *A*^*m*^ of a network *m*, a traditional GAT computes an embedding for a node by aggregating the features of its neighboring nodes but after giving different priorities or “attention” values to each neighbor. We compute the attention 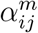 that node *i* gives to node *j* in network *m* as follows:

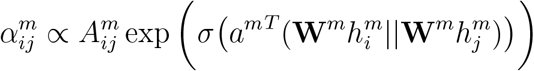

Here,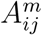is 1 iff *i* and *j* are connected by an edge in network *m*, 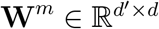 is a network-specific learnable weight matrix where *d* and *d*^′^ are input and output feature dimensions respectively for the GAT layer, 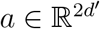 is a vector of learnable attention coefficients. *σ* is a non-linear activation function, e.g., LeakyRelu, *h*_*i*_ is the feature vector of node *i* that is input to the GAT layer, and || denotes concatenation operation. The constant of proportionality in the equation ensures that the sum of 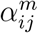 over all the neighbors of node *i* equals 1.

#### Co-attention GAT

For network *m*, we compute an *n* × *n* co-attention matrix *χ*^*m*^, which is a weighted average of the already computed attention from all the input networks in the previous GAT layers. Specifically, we compute 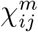, the co-attention that node *i* gives to node *j* as follows:

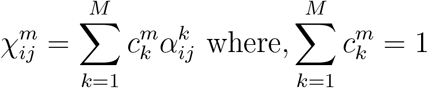

Here, *M* is the total number of input networks and the learnable *co-attention coefficient*, 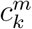 denotes the priority given by network *m* to (the attention computed by the GAT layer on) network *k*.

Finally, we compute the embedding for node *i* in network *m* as follows:

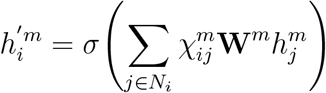

This new vector will be the input for the next GAT layer.

#### Network reconstruction module

The encoder module generates separate embeddings of a protein for each of the input networks. The first step of the reconstruction module is to generate a unified embedding for each protein by taking an average of its embeddings across all the networks. To construct the integrated network, we utilize a simple dot product operation on the unified features of pairs of nodes. Suppose, matrix *F* contains the unified embeddings of each node in the network. We compute the adjacency matrix *Â* of the reconstructed network as follows: *Â* = *F*.*F*^*T*^

#### Loss computation

We train ICoN to minimize the discrepancy between the reconstructed network and the original input networks (without noise). This strategy empowers our model to navigate through noisy networks to reach a denoised integrated network. We formulate the network reconstruction loss *L* as follows, where *n* is the total number of nodes and ||.||_*F*_ is the Frobenius norm: 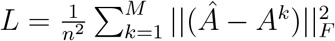

### 2.2 Datasets

We used the networks and evaluation strategy created by BIONIC [4] and subsequently utilized by BERTWalk [9]. We briefly describe this framework in this section and the next. We refer the reader to the BIONIC publication for a detailed description.

We ran ICoN on heterogeneous networks originating from diverse experiments for *Saccharomyces cerevisiae* and human. We integrated three baker’s yeast (i.e., *Saccharomyces cerevisiae*) networks: (i) a protein-protein interaction (PPI) network (2, 674 genes and 7, 075 interactions) obtained by tandem affinity purification followed by mass spectrometry [15], (ii) a genetic interaction (GI) network (4, 529 genes and 33, 056 interactions) constructed by calculating the pairwise Pearson correlation between genetic interaction profiles of genes [1], and (iii) a coexpression (COEX) network (1, 101 genes and 14, 826 interactions) based on the Pearson correlation between transcriptional response profiles of deletion yeast strains [16]. We also integrated four human protein-protein interaction networks emerging from diverse experimental approaches: (i) Rolland-14 (4, 301 genes and 13, 940 interactions) [17], (ii) Hein-15 (5, 380 genes and 27, 349 interactions) [18], (iii) Huttlin-15 (7, 658 genes and 23, 712 interactions) [19], and (iv) Huttlin-17 (10, 945 genes and 56, 471 interactions) [20]. This analysis permitted us to study ICoN’s generalizability across species and determine if there was any utility in integrating networks containing the same type of interactions (PPIs) but generated by different groups using complementary experimental techniques.

### 2.3 Evaluation

The evaluation framework consists of three downstream tasks. The ground truth for evaluating performance on those tasks is extracted from three benchmark datasets.

#### Downstream tasks

The three tasks are the following:

i. gene coannotation prediction: The goal of this task is to evaluate how well a model preserves gene-gene relationships in its node embeddings. We predict whether a pair of genes are annotated to the same term in a particular functional benchmark. Given the embeddings for a pair of genes we compute the cosine similarity. We categorize gene pairs as positive (i.e., coannotated) or negative based on a predefined threshold. By varying this threshold, we compute the average precision.
ii. gene module detection: Here we assess the model’s ability to reproduce biological modules such as protein complexes, pathways, and biological processes by performing hierarchical clustering of nodes (based on embeddings) using a variety of distance metrics (Euclidean, cosine) and linkage methods (single, average, and complete). To evaluate the similarity between the derived clusters and the ground truth biological modules (from a specific benchmark), we calculate the adjusted mutual information (AMI) score. We optimize the clustering parameters, (i.e., distance metric and linkage method) for each network integration method (and network) and report the best AMI.
iii. gene function prediction: This task seeks to evaluate a model’s capability to generate embeddings that are effective in predicting known functional classes (such as membership to a particular protein complex, biological process, or pathway). This task entails training a support vector machine classifier using node embeddings as features to predict gene functions.

#### Benchmarks

We used four datasets (the first three for yeast and the final one for human): (i) IntAct protein complexes [21], (ii) Kyoto Encyclopedia of Genes and Genomes (KEGG) pathways [22], (iii) Gene Ontology biological processes (GO BP) [23], and (iv) CORUM complexes [24]. In each of these datasets, genes are annotated to functional terms indicating their membership in certain protein complexes, pathways, or biological processes. To create a coannotation benchmark from any of these datasets, all gene pairs that share at least one annotation are considered to be positive pairs, while those lacking shared annotations are classified as negative pairs. The coannotation datasets for IntAct, KEGG, and GO BP contained 1, 786, 2, 029, and 4, 170 genes and 9, 288, 32, 563, and 45, 015 positive coannotations, respectively. We constructed the benchmarks for module detection by defining a module as a collection of genes annotated to the same functional term whereas the benchmark for function prediction was formed by considering each gene annotation as a class label. The module detection benchmarks included 574, 107, and 1, 809 modules and the gene function standards consisted of 48, 53, and 63 functional classes, respectively for IntAct, KEGG, and GO BP. Finally, the human CORUM dataset contained 3, 431 genes and 39, 011 positive coannotations.

### 2.4 Hyperparameter optimization

To optimize hyperparameters for ICoN we used the same approach as BIONIC. We ran ICoN on three *S. pombe* networks, i.e., a genetic interaction network [25], a coexpression network [26], and a protein–protein interaction network [27]. We evaluated the embeddings generated by different combinations of hyperparameters on gene module detection, coannotation prediction, and function prediction tasks where we utilized an *S. pombe* protein complex dataset as a gold standard.

We fine-tuned two hyperparameters: the number of layers in the encoder module (2 or 3) and the noise induction rate (0.1, 0.3, 0.5, and 0.7). ICoN with 2 encoder modules and noise induction rate of 0.7 achieved the highest average rank across the three tasks. We used these hyperparameters of ICoN henceforth. We ran BIONIC [4], deepNF [8], and Mashup [7] with the best hyperparameters reported by BIONIC. Since the BERTWalk paper [9] does not mention hyperparameter optimization, we evaluated it on the author-provided embeddings.

## 3 Results

First, we assessed the performance of the integrated features obtained through ICoN against features from individual input networks, based on different downstream tasks across varied functional benchmarks (Section 3.1). Next, we performed a comparative analysis of ICoN with four unsupervised network integration methods: Mashup [7], deepNF [8], BIONIC [4], and BERTWalk [9] (Section 3.2). We ran all algorithms five times, except for BERTWalk. Third, through an ablation study, we showcased the utility of two pivotal components of ICoN: co-attention GAT and noise induction (Section 3.3). Subsequently, we investigated the learned co-attention weights (Section 3.4). Finally, we analyzed the robustness of ICoN to noise in the input networks (Section 3.5).

### 3.1 Improvement over individual input networks

In our first analysis, we observed that the embeddings generated by ICoN outperformed the features from each of the individual yeast networks in all three tasks across all three functional benchmarks (Figure 2a). Here for individual input networks, we regard the interaction profile of proteins as features. The same observation held for the four human PPI networks in the coannotation prediction task in the CORUM complexes benchmark (Figure 2b). Note that the absolute performance of ICoN-based features changes from task to task and from one benchmark to another, as indicated by the different ranges of the *y*-axes in the plots (Figure 2). Nevertheless, these results confirmed the efficacy of employing integrated features as opposed to relying on individual networks.

**Figure 2:**
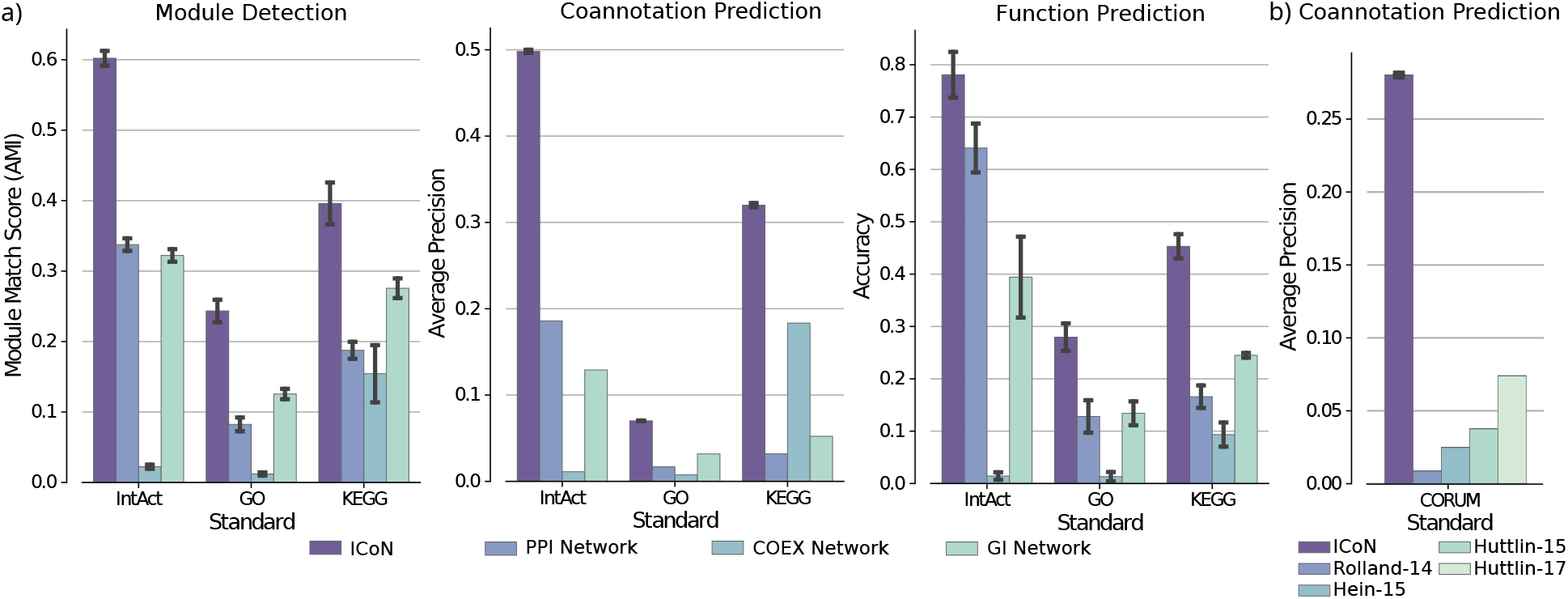
Evaluate ICoN’s performance on downstream tasks. Compare the performance of ICoN with a) three yeast networks and b) four human networks.

### 3.2 Improvement over existing unsupervised methods

We compared ICoN to BERTWalk [9], BIONIC [4], deepNF [8], and Mashup [7] and to Union, a naive approach where we computed the integrated network as the union of the node sets and edge sets across all the input networks.

For the three yeast networks, ICoN outperformed all the methods in module detection across all three functional benchmarks (Figure 3a). In each of these benchmarks, either BIONIC or BERTWalk emerged as the second-best model, with no single model consistently maintaining the second position. ICoN obtained a significantly higher AMI compared to the second-best model for all three benchmarks (Mann-Whitney U test *p*-values of 1.4 × 10^−13^ in IntAct, 0.01 in GO, and 0.0015 in KEGG). We also observed that ICoN surpassed each of the models in coannotation prediction in IntAct and GO benchmarks showing a significantly higher average precision score than the second-best model BIONIC (Mann-Whitney U test *p*-values of 0.006 in both benchmarks) while showing comparable performance to BIONIC on the KEGG dataset. BERTWalk outperformed ICoN in the KEGG benchmark for the co-annotation task. We could not estimate the statistical significance of this difference since we could run BERTWalk only once (with the author-provided embeddings). Finally, in function prediction, ICoN exhibited comparable performance, with no statistically significant difference, to the best-performing model, BIONIC, in two out of three functional benchmarks (i.e., IntAct and GO). However, in the KEGG benchmark, ICoN demonstrated inferior performance compared to BIONIC (Mann-Whitney U test *p*-value 4.43 × 10^−5^). For the human datasets, ICoN demonstrated a persistent pattern of excellence by outranking all five methods (with Mann-Whitney U test *p*-value 0.006 for performance differences between ICoN and BIONIC) (Figure 3b).

**Figure 3:**
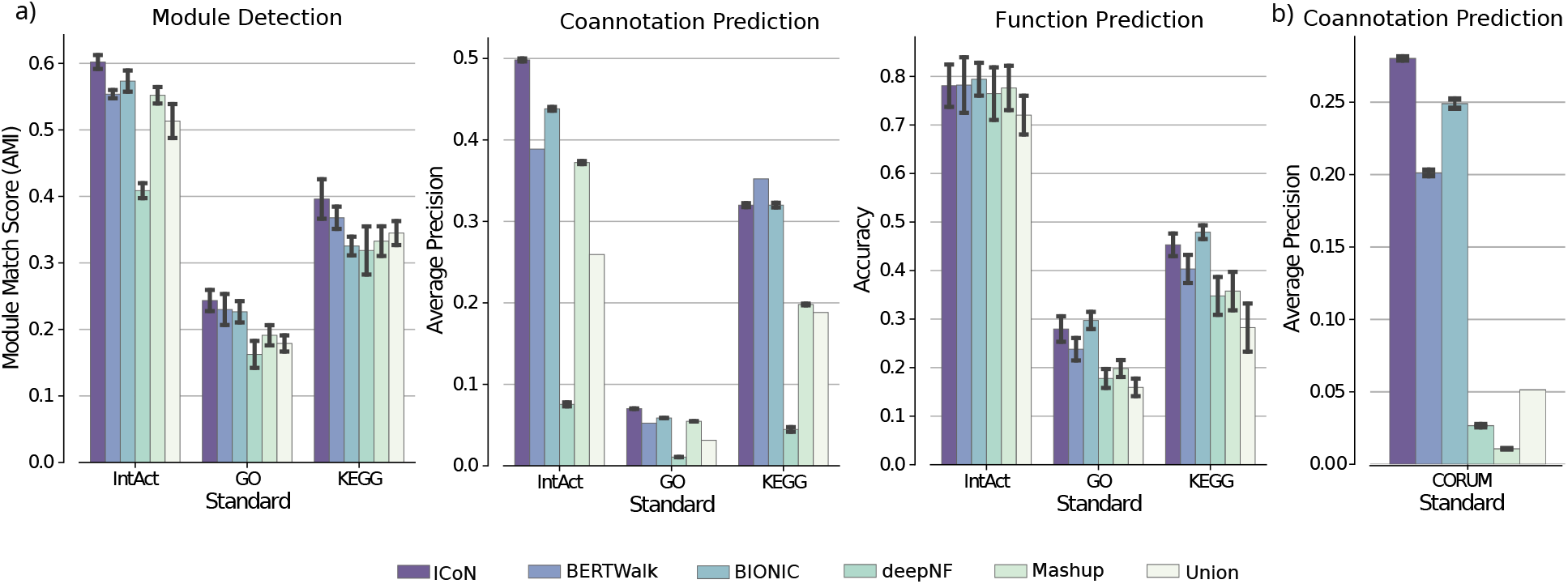
Comparison of ICoN and five unsupervised network integration methods on a) three yeast networks and b) four human networks.

In summary, ICoN outperformed all the methods in six out of ten task-benchmark pairs while showing comparable performance to the best-performing model in two. Additionally, it is noteworthy that in the two tasks where ICoN had the second-best performance, no single model emerged as the indisputable best performer.

### 3.3 Ablation study

We introduced co-attention in ICoN to facilitate the exchange of topological information across networks during training. To study the importance of co-attention, we conducted an experiment where we constrained ICoN to solely consider self-computed attention to aggregate features from its neighbors, thereby eliminating co-attention. We found that ICoN (i.e., with co-attention) outperformed the restricted model on the majority of downstream tasks (six out of nine with Mann Whitney U test *p*-value *<* 0.007) (Figure 4a).

**Figure 4:**
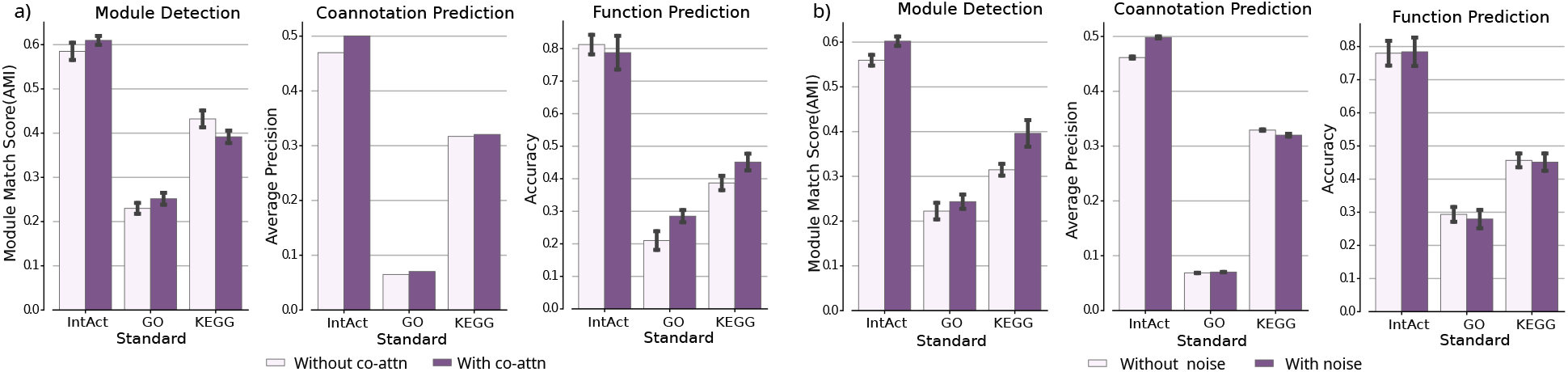
Results of ablation study on co-attention and noise induction.

In the second ablation study, we avoided the induction of noise into the input networks. Our findings indicate that the absence of noise adversely affects performance in the majority of downstream tasks (Mann Whitney U test *p*-value *<* 0.007) (Figure 4b).

It is noteworthy that either co-attention or noise induction helps in every downstream task. Their complementarity enables ICoN to surpass existing network integration models.

### 3.4 Interpretation of the learned co-attention coefficients

Here, we aimed to assess to what extent ICoN exploited its ability to share attention (i.e., co-attention) across networks. Additionally, we sought to investigate whether one network can assist another in mitigating the impact of noise through this shared attention mechanism. We first observed that for every value of noise that we tested, each of the three yeast networks learned non-zero co-attention coefficients not for just itself but for other input networks as well (Figure 5). For most values of noise, a network gave the highest priority (i.e., a co-attention coefficient) to itself. As we increased noise, the priority given by the GI network to itself decreased substantially: from 0.69 at 0% noise to 0.36 at 50% noise (Figure 5a). The COEX network also showed this pattern of lowered self-priority with heightened noise, although the trend was not as marked as for the GI network. In contrast, the learned co-attention coefficient by the PPI network for itself increased slightly with noise: from 0.47 at 0% noise to 0.53 at 50% noise. We also found that as we increased noise, the PPI network was given more priority by the other two networks: from 0.38 at 0% noise to 0.67 at 50% noise (Figure 5b).

**Figure 5:**
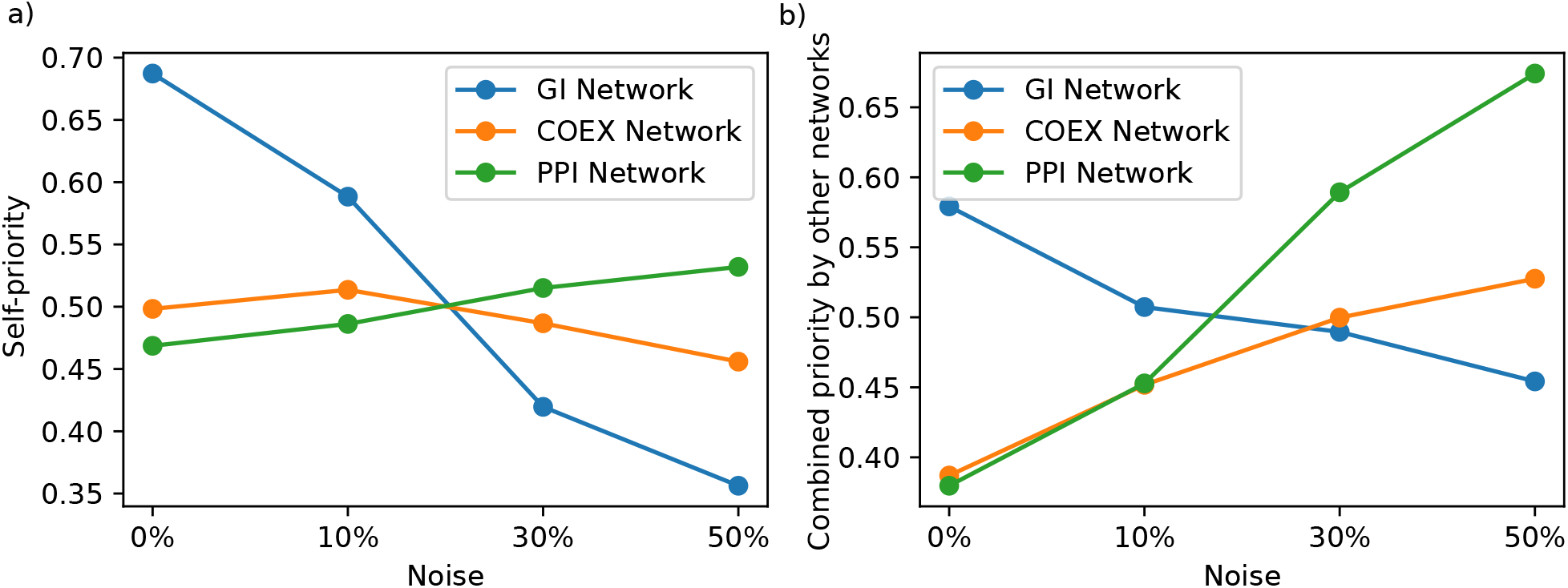
Variation in co-attention coefficient with noise (shown for the first layer of encoder module). Along the *x*-axis, we show the amount of noise introduced by the noise induction module. Along the *y*-axis we show a) the amount of self-priority (i.e., learned co-attention coefficient) each network gives to itself and b) the total amount of priority each network gets from the other networks.

The GI network is the largest, encompassing 60% of the total number of edges, while the COEX and PPI networks contain 27% and 13% of the edges, respectively (Section 2.2). The larger the network, the more the number of true edges removed or false interactions introduced by noise induction. We surmised from Figure 5 that the two largest networks mitigated the impact of this noise by assigning themselves reduced self-priority. In contrast, the PPI network refrained from adopting this approach, as diminishing self-priority would mean giving increased attention to the other networks, resulting in a higher influence of noise from them. This explanation also accounts for the increased amount of priority allocated to the PPI network by the others as noise escalates.

### 3.5 Robustness to noise

A proposed network integration method should be robust to the presence of noise. To assess this property, we artificially introduced noise by dropping a certain percent of existing edges and then adding the same number of random edges to each of the original input networks. Then we integrated the noisy networks employing ICoN, BIONIC, and Union. Unlike noise induction, where we introduce noise into the input networks and minimize the reconstruction loss on the original edges, here we measure the loss with respect to the noisy network.

In this analysis ICoN maintained its superiority in module detection against BIONIC as the networks became noisier (Figure 6a) demonstrating ICoN’s robustness in handling noisy input networks. It is also noteworthy that, while ICoN retained 81% and 60% of its AMI score as the noise increased to 30% and 50% respectively, BIONIC only retained 74% and 53% of its original AMI (Figure 6b).

**Figure 6:**
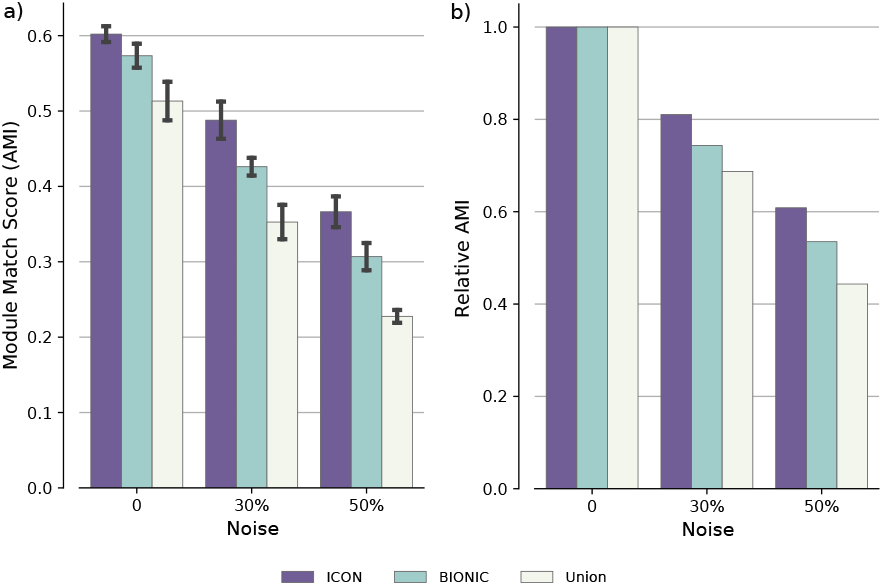
Comparison of ICoN, BIONIC, and Union in robustness to noisy inputs.

## 4 Conclusions

In ICoN, we introduce a novel graph neural network architecture that facilitates attention sharing across networks to generate unified, integrated protein embeddings. This unsupervised model, trained solely on network topology, exhibits superior generalization across the majority of downstream tasks compared to existing biological network integration models. Furthermore, ICoN demonstrates robustness against noisy networks. A potential future avenue involves incorporating gene or protein features such as sequence and structure data, along with protein embeddings from pretrained models.

